# Abstract choice representations during stable choice-response associations

**DOI:** 10.1101/2024.10.17.618830

**Authors:** Katrina R. Quinn, Florian Sandhaeger, Nima Noury, Ema Žeželić, Markus Siegel

## Abstract

Perceptual decisions have long been framed in terms of the actions used to report a choice. Accordingly, studies of perceptual decision-making have historically relied on tasks with fixed choice-response mappings, in which choice and motor response are inextricably linked. Although several studies have since dissociated choice and response, they have typically involved dynamic switching of the choice-response mapping on a trial-by-trial basis. Thus, it remains unclear if abstract choice representations arise specifically when choice-response relationships change dynamically, or if they reflect a more general property of the decision-making process. Here, we show that in the human brain, choices are represented abstractly, even when the association between choice and motor response remains stable over time. We measured neural activity in humans using magnetoencephalography (MEG) while participants performed a motion discrimination task. Importantly, the associations between perceptual choice and motor response were balanced across experimental conditions and remained stable over many trials. We found neural information about the participants’ perceptual choice, independent of both the motor response and the visual stimulus. This abstract choice information increased during the stimulus period and peaked after the response had been made. Furthermore, choice and response information showed distinct cortical distributions, with strongest choice information in frontoparietal regions. Our results suggest that abstract choice representations are not restricted to action-independent contexts or those with dynamic choice-response associations and may therefore reflect a general role in perceptual decision-making.

## Introduction

Many of our every-day decisions are tightly coupled to a particular motor action. Likewise, most of the tasks exploring the neural underpinnings of perceptual decision-making have inextricably linked choices with the motor-responses required to report them [1–5], for example in motion discrimination, a rightward eye movement or button press for a downwards motion choice. This is in keeping with one of the dominant accounts of perceptual decision-making, which suggests that choices are embodied [6–8]. This intentional framework suggests that choices emerge as plans to commit a particular action.

However, we are also able to make decisions that cannot immediately be followed by a specific action. This implies that the brain can represent choices independently of actions, which we here refer to as abstract choices. How this is achieved, and under what contexts such representations may persist, has been a matter of continued debate [2,3,6,8– 10]. In recent years, there has been a greater focus on uncoupling choices and actions in perceptual tasks, to identify potentially independent neural representations. Indeed, these studies have found choice signals which arise before the choice-response mapping is known, therefore abstracting them from any representation of the motor-response [11– 15]. More importantly, the same abstract choice signals were identified regardless of whether the choice-response mapping was known in advance or not [15], indicative of an abstract choice stage across different action contexts.

The extent to which abstract representations of choice occur ubiquitously across different task contexts remains unclear. One possibility is that an abstract choice stage is specifically recruited when flexible shifting between choice-action mappings is required. Previous studies separating choice and action representations have typically used either post-stimulus (action-independent) mapping [12–14,16,17] or trial-by-trial switching either between action-independent and action-linked decisions [15] or between different pre-stimulus choice-action mappings [18]. Alternatively, an abstract choice stage may play a general role in decision-making, even when little to no flexibility of choice-action associations is required.

We investigated these alternatives in the human brain by holding the choice-response mapping stable over extended periods of time. Human participants performed an up-down visual motion discrimination task while we recorded neural activity using magnetoencephalography (MEG). Importantly, using a blocked task design and specific analyses, we were able to disentangle choice and response representations. We found neural activity selective for the stimulus, motor response and choice. Crucially, choice information was independent of the choice-response mapping. This abstract choice representation ramped up during the stimulus period, consistent with evidence integration over the motion stimulus, and peaked after the response. Our results show that choices are represented abstractly, even when the association between choice and motor response remains stable over time. This suggests that abstract choice representations may play a more general role than purely action-based frameworks have previously implied.

## Results

### Task and behaviour

We recorded MEG in 16 human participants (20 prior to exclusions; see Methods) while they performed an up-down visual motion discrimination task (Fig 1A). On each trial, the participants viewed a stream of 24 random-dot motion pulses (83 ms per pulse). After an auditory go-cue they indicated whether they perceived more up- or downward motion with a left or right button press. The mapping between choice (up or down) and response (left or right button) was manipulated block-wise, such that all trials within a single block of 96 trials had the same choice-response mapping (e.g. left button press for upward motion, right for downward). The mapping was indicated to the participants at the beginning of each block. The mapping for each block was determined pseudo-randomly, such that mappings were balanced across stimulus conditions (see below). Most often, this meant that the choice-response mapping switched between blocks but could also remain stable over multiple blocks (Fig 1B).

**Figure 1.**
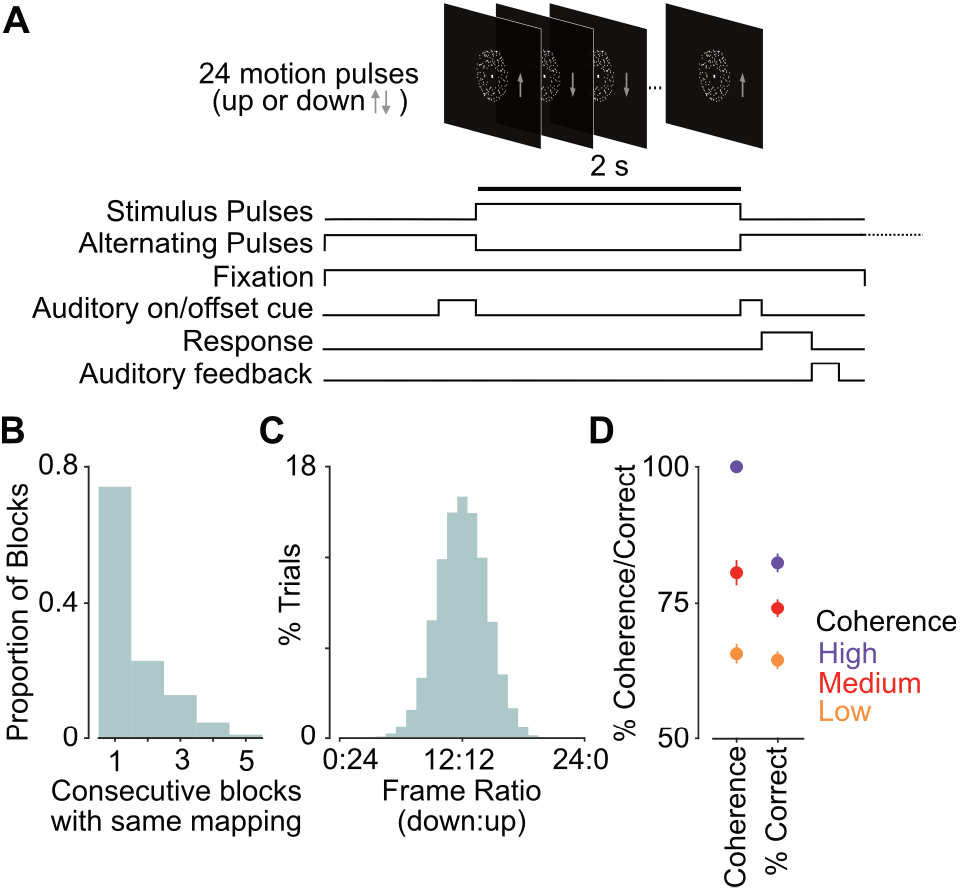
Up-Down motion discrimination task and behavioural performance. (**A**) On each trial, participants (n = 16) were required to discriminate the average motion direction of 24 random-dot motion pulses. Choice was indicated with a left or right button press. The mapping of choice to response was pseudo-randomised block-wise (96 trials per block), such that in one block a left button press corresponded to an ‘up’ choice, and in another block to a ‘down’ choice. Grey arrows depict average pulse motion, shown here for visualisation purposes only. (**B**) Histogram of the proportion of successive blocks with stable choice-response mapping. (**C**) Ratio of ‘up’ to ‘down’ motion pulses as percentage of trials. Each pulse was drawn randomly with equal probability. 12:12 corresponded to an equal number of ‘up’ and ‘down’ pulses on a trial, for which participants were rewarded randomly. (**D**) Motion coherence and correct performance for the three coherence levels (low, medium, high). Coherence was pseudo-randomly varied across blocks. Low and medium coherence levels were adjusted for each participant to target 66% and 75% correct performance, respectively, using staircases prior to the task. High coherence was 100%. Error bars denote SEM across subjects.

Each of the 24 motion pulses showed either upward or downward motion. The motion direction was determined randomly on a pulse-by-pulse basis, such that each individual pulse had an equal probability of being an up or downward motion pulse (Fig 1C). The strength of motion for each pulse was determined by the motion coherence, which was held constant within a block. There were three coherence levels in total – low, medium and high. Low and medium coherence levels were determined by staircases prior to the task to achieve about 66% and 75% performance, respectively. In the high coherence condition, 100% of the dots moved in the specified direction. The order of the coherence blocks was determined pseudo-randomly, such that the coherence levels were balanced.

The coherence staircases had the desired effect that the three coherence levels were well-separated (Fig 1D, left; mean coherence: 66%, 81%, 100% for low, medium and high coherence, respectively). Behavioral performance corresponded well with the expected performance for the respective coherence levels (Fig 1D, right; mean correct performance: 65%, 74%, 82% for low, medium and high coherence respectively).

### Neural information about motion direction and motor responses

We quantified neural information about the direction of motion pulses and about motor responses for each participant using a cross-validated multivariate analysis of variance (cvMANOVA) [15,19,20]. We applied cvMANOVA on preprocessed MEG data across all sensors (see Methods), resulting in a measure of neural information analogous to classifier performance. Importantly, cvMANOVA independently assesses the variability related to a specific variable of interest, while excluding confounds related to other variables.

We were able to quantify neural information about individual motion pulses, which can be observed as the shifting peaks of information across time relative to stimulus onset (Figure 2A). When we aligned individual pulses by their onset time and averaged neural information across pulses, we found significant information for each coherence level (Fig 2B; p = 0.03, 0.02, 0.017 for low, medium and high coherence, respectively, cluster permutation statistics). Motion information peaked about 300 ms after pulse onset and continued for several hundred milliseconds. Information about motion direction increased with increasing motion coherence, although this effect was not statistically significant (F = 1.23, p = 0.30, one-way ANOVA of average pulse information 200-400 ms post pulse onset with factor coherence). As the cvMANOVA also included the subjects’ choice and motor response as factors, neural information about motion pulses was independent of these factors.

**Figure 2.**
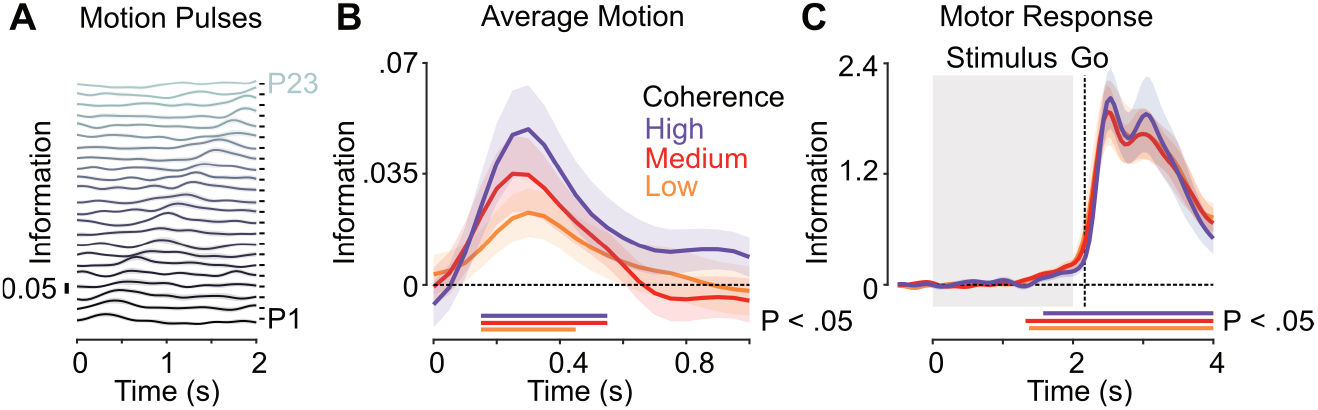
Neural information about motion direction and motor response. (**A**) Neural information about the direction of each motion pulse averaged across coherence levels. P1 and P23 denote the first and second-to-last pulse respectively. Time axis spans stimulus onset to offset. (**B**) Neural information averaged across pulses 1-23, aligned to pulse onset, for each coherence level. (**C**) Neural information about the side on which the button was pressed (left vs. right) for each coherence level. Horizontal bars denote temporal clusters of significant information (p < 0.05; cluster permutation). Solid lines and shaded regions indicate the mean +/- SEM of information across participants (N = 16).

We also found significant neural information about motor responses for each coherence level (Fig 2C; p = 0.002, 0.003, <0.001 for low, medium and high coherence, respectively; cluster permutation). Again, as the cvMANOVA also included the subjects’ choice as a factor, neural information about the response was independent of the choice. Response information started to rise in the second half of the stimulus period, consistent with early motor response preparation, and peaked after the go-cue. We did not observe any differences in the response information as a function of coherence (F = 0.01, p = 0.99, one-way ANOVA of average response information 0-1s post stimulus offset with factor coherence). In summary, we found robust neural information about the sensory stimulus that subjects had to decide upon and about the motor response with which subjects reported their choice.

### Neural information about choices independent of responses

We next tested if we could find neural information for the perceptual choice independent of the motor response and stimulus. To do so, we first corrected neural activity for any influence of the stimulus, so that neural activity related to choice would not be confounded by the correlations which arise between stimulus and choice for above-chance performance. We implemented a pulse-based stimulus correction because, unlike for the response or choice variable, merely including the average stimulus motion as a factor in the cvMANOVA would not render other information independent of stimuli at the pulse level (see Methods). Critically, the employed correction was conservative, i.e. would lead to an underestimation of choice information dependent on the behavioural performance. As performance differed between coherence levels, we restricted the subsequent analysis to the data averaged across coherence conditions. Furthermore, as the cvMANOVA also included the response as a factor, choice information was assessed independently from the motor response. In other words, we assessed abstract choice information.

We found significant neural information about the perceptual choice (Fig 3A; p = 0.001, cluster permutation) which ramped up during the stimulus period (p = 0.015, one-tailed t-test for the stimulus presentation interval) consistent with evidence integration over the motion pulses and peaked after the go-cue. When we aligned the analysis to the response time based on the reaction time on each trial (Fig 3B, p=.001, cluster permutation), we found that choice information largely peaked right after the response and declined thereafter.

**Figure 3.**
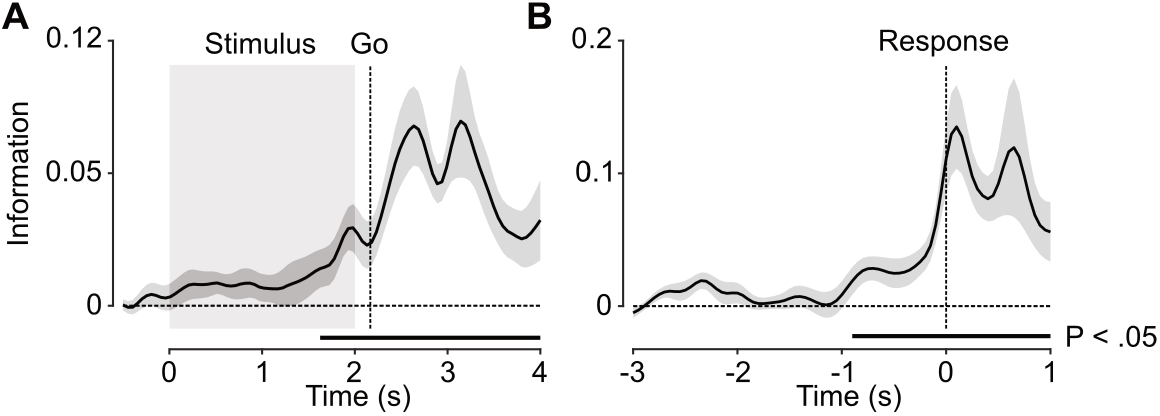
Neural information about choice. (**A**) Neural information about the participants choices (up vs. down) averaged across coherence conditions and aligned to stimulus onset (n = 16). (**B**) Choice information aligned to the time of response (n = 14). Horizontal bars denote temporal clusters of significant information (p < 0.05; cluster permutation). Solid lines and shaded regions indicate the mean +/- SEM of information across participants.

### Choice and response information exhibit distinct cortical distributions

To further contrast choice and response information, we employed a searchlight analysis and quantified their cortical distribution across time, averaged across hemispheres (see Methods). We found distinct patterns of information both across space and time (Figure 4). For response information, we observed a central peak after the go-cue which continued well into the response period. In contrast, choice information was strongest in parietal and frontal regions during the stimulus period. This pattern continued into the response period, with additional central contributions.

**Figure 4.**
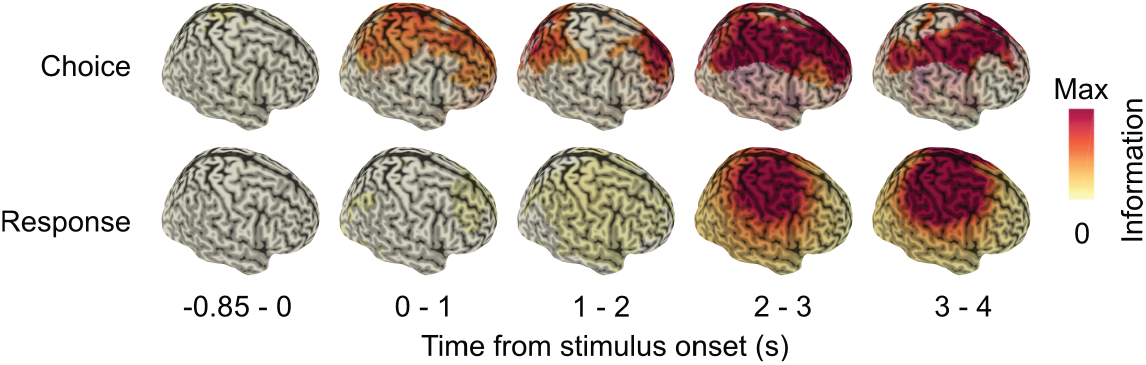
Cortical distribution of choice and response information. The spatiotemporal distribution of choice (upper row) and response (bottom row) information for 5 temporal intervals, with respect to stimulus onset (0 s). Only significant clusters of information are shown (p < 0.05; cluster permutation).

### Choice information includes response-independent and response-linked components

Our experimental design and analysis approach ensured that the identified choice information was independent of the motor response, i.e. that the identified neural variability that was explained by choices could not be explained by motor responses. However, choice and response information may still recruit overlapping neural populations. To test this, we quantified the cross-variable information between choice and response (see Methods). We found that the absolute choice-response cross-variable information was significantly lower than expected if they involved perfectly overlapping neural populations (Fig 5A; paired t-test, p = 0.000027). However, absolute cross-variable information was also significantly above chance, indicating that the representations of choice and response were not completely orthogonal (Fig 5B; one-tailed paired t-test, p = 0.00026 real vs. shuffled choice labels). This suggests that choice and response information arise from neural populations that are largely disparate but may have a small degree of overlap.

**Figure 5.**
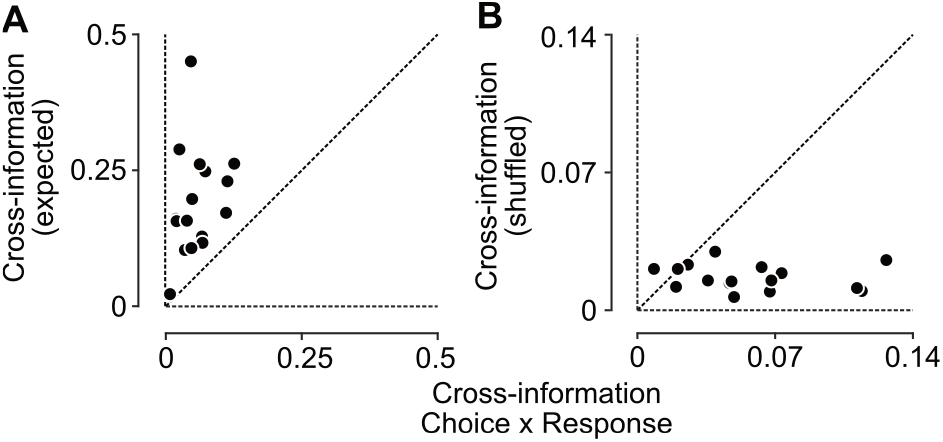
Cross-variable information between choice and response. (**A**) Absolute cross-variable information is significantly lower than expected for identical neural patterns of choice and response information. (**B**) Absolute cross-variable information between choice and response was significantly higher than expected by chance for data with shuffled choice labels. Each dot represents a single participant. All analyses are for the time interval 1.5-3 s post stimulus onset.

## Discussion

We found neural information about perceptual choices independently of motor responses in the human brain. These results are compatible with previous findings showing abstract choice representations both when the choice-motor mapping is known in advance [15,18], and also when it is not [11–17]. Importantly, we found abstract choice information in a task where the choice-response mappings remained stable over many trials. This contrasts with the previously mentioned studies where choice-motor mappings changed on a trial-by-trial basis. Thus, our results are, to the best of our knowledge, the first to show that abstract choice representations are not limited to tasks which require a quick and flexible mapping of choices onto responses and may therefore reflect a more general property of the perceptual decision-making process.

Our results complement a recent study which used similar methods to show that abstract choice information generalised between action-linked and not action-linked decisions. The present study provides two main advances. Firstly, as mentioned above Sandhaeger et al. [15] changed the choice-motor mapping and when it was cued on a trial-by-trial basis, whereas in our study the mapping was always known in advance and held stable across many trials. Secondly, Sandhaeger et al. used an asymmetric task design, where participants decided if coherent motion was present or not. Using a symmetric discrimination in the present task (up vs. down) ensures that the variables of interest are less likely to be affected by other extraneous variables, such as e.g. arousal. Specifically, for any extraneous variable to explain choice information, it would need to act differently between trials for the two choices, which is unlikely when the choice options are symmetric, e.g. around the axis of motion.

We found that choice information was present in a network of brain areas distinct from that for the motor response, with parietal and frontal areas showing significant choice signals. These results are broadly consistent with previous findings in non-human primates [11] and humans [12,15,16,21]. They also complement a recent study which localised an indirect measure of choice activity to fronto-parietal networks when motor-responses are held stable [22]. Previous evidence in humans has shown conflicting results on the involvement of posterior parietal cortex, with some suggesting that this area only play a role in action-linked contexts. Although our task involves action-linked decisions, our results show choice-related activity in a parietal region, independent of the motor response. One explanation suggested by Filimon et al. [16] could be that posterior parietal cortex is involved in representing abstract choices in a motion discrimination task, due to its strong connections with area MT, but not in other perceptual discriminations. Alternatively, posterior parietal cortex may multiplex signals associated with choice and response [23], in such a way that it is not always clear from human brain imaging whether these regions are involved in the task at hand. Further studies, using invasive recordings or more complex experimental designs are required to elucidate this issue.

The fact that we found abstract choice representations even when the choice-response association was stable over time suggests that these representations are more ubiquitous than previously thought. The intentional framework of decision-making has long viewed representations of choice to be synonymous with convergence upon an action-plan, framing abstract decision contexts almost as outliers [6,8]. While there has been a wave of evidence for abstract choice representations when the choice-response mapping is not known in advance of the stimulus, only recently has there been evidence to suggest that these persist even when the mapping is known in advance [15,18]. Our findings contribute to this evidence. Furthermore, our results address a criticism of abstract choice tasks, that participants may still plan response actions even when the mapping is not known in advance [24]. If individuals preferentially associate a specific choice with a specific motor response, this could mean that evidence for abstract choice representations is in fact still motor-linked. However, our cross-variable information analysis shows that even on an individual level, the neural populations supporting choice and motor-response are largely distinct. Together, these findings contradict a purely intentional framework, and at the very least suggests that abstract representations of choice are not limited to contexts in which an action cannot immediately be planned.

Our results lead us to several open questions. What is the precise role for abstract choice representations when an action can be planned directly? A recent study from Charlton & Goris (2024) found that information about the choice emerged earlier than that about the response, suggestive of a choice stage prior to the action planning. In our study, choice and response information emerged at similar times, although the different strength of the signals complicates direct timing comparisons. In any case, even without a serial processing of choice and action plan, it could be beneficial to represent choices in an abstract reference frame. For example, when learning to navigate our natural environment, the coupling between particular choices and actions varies with changes of our viewpoint or dynamic properties of the visual scene. In this case, it could be advantageous to know the perceptual choice associated with particular visual stimuli and outcomes, in addition to the latter’s association with specific actions.

In which contexts do these abstract choice representations arise? In our study it was methodologically necessary to change the choice-response mapping several times to measure choice information independently of the visual stimulus and the motor response. While the blocked design was more stable than a trial-by-trial design, it could still be that the infrequent changes of the choice-response mapping led to choices being represented in an abstract format generalising across block types. This leaves open the possibility that in contexts where the choice and associated action are even more tightly coupled, decision-making operates in a fully intentional way, without abstract choice representations. However, given that our natural behaviour involves many instances of both abstract and action-linked perceptual decisions, it seems intuitive to implement neural processes that allow to fluidly switch between these different contexts. To this end, choices could be represented abstractly across all contexts, potentially simultaneously with action-linked choice signals, and utilised when required.

## Methods

### Participants

20 healthy, right-handed human participants (5 female; mean = 27 years, 4 year SD) took part in the current study and received monetary reward. All participants had normal or corrected-to-normal vision. Prior to the recording, participants provided written informed consent. The study was approved by the ethical committee of the Medical Faculty and University Hospital of the University of Tübingen and conducted in accordance with the Declaration of Helsinki.

### Behavioural task and stimuli

Participants performed a symmetric motion discrimination task. On each trial, they were asked to report whether a train of 24 motion pulses contained more upwards or downwards motion. Responses were made with a left- or right-hand button press. Importantly the mapping between choice and motor response varied block-wise, such that a left button press would correspond to an up or down choice depending on the block.

At the start of each trial participants were required to fixate a small white point at the centre of the screen, and continued fixating for the duration of the trial. Once they obtained fixation, 16 alternating up/down motion pulses (1/12 = 0.08333 s pulse duration) were presented (1.33 s total duration), centred around fixation. For the last 0.3333 s of this pre-stimulus pulse train, an auditory onset cue signalled the start of the stimulus pulses. Subsequently 24 stimulus pulses (1/12 = 0.08333 s pulse duration) were presented (2 s total stimulus duration), each showing more up or downward motion. After the stimulus pulses, a post-stimulus pulse train of alternating up/down pulses began. At the same time, an auditory offset cue (0.1667 s duration) signalled the end of the stimulus-train and acted as a go-cue for participants to respond. After the response, participants received auditory feedback about their performance. Following the feedback, a random even number of alternating up-down pulses (4-14 pulses) was presented before the next trial began.

Participants sat at a viewing distance of 50 cm from the screen. Fixation occurred within a window of 2 degrees (7 subjects) or 1.34 degrees (9 participants) of visual angle. The fixation dot was a white dot with a radius of 0.1 degrees. A single motion pulse consisted of a random-dot kinematogram with 700 white dots, each with a radius of 0.1 degrees. The dots were presented within a circular aperture on a black background (6.7 degrees radius), with an additional circular aperture without dots surrounding the fixation point (1.34 degrees radius). Each dot moved either upward or downward at 10 degrees per s. The proportion of dots moving upwards vs. downwards was determined by the motion coherence of which there were three levels – low, medium and high. For the high level, all dots moved in one direction. Low and medium coherence levels were determined using two 2:1 staircases (144 trials each) prior to each experiment, to achieve about 66% and 75% correct motion discrimination performance, respectively. For each coherence level, there were 10 possible ‘up’ pulses, which were flipped to produce 10 possible ‘down’ pulses. During the experiment, for each of the 24 stimulus pulses, the pulse direction and specific pulse (1 of 10) was drawn randomly with equal probability. For stimuli with the same number of ‘up’ and ‘down’ pulses (12:12), subjects received randomized feedback.

The auditory onset and offset tones had a frequency of 200 and 300 Hz, respectively. The auditory feedback consisted of two tones: one linearly increasing from 200 to 600 Hz and the other in the reverse direction. The mapping of the two tones to feedback (i.e. correct or incorrect) was varied blockwise.

Each block consisted of 96 trials. Participants typically completed 4 repetitions of each of the 3 types of coherence level blocks, totalling 12 blocks. Each repetition included one block at each coherence level. The choice-motor mapping, auditory feedback mapping and coherence level for each block were determined pseudo-randomly, and participants were informed of the choice-motor and feedback mappings at the beginning of each block.

### Setup and recording

Neural activity was recorded using a 275-channel whole-head MEG system (Omega 2000, CTF Systems, Port Coquitlam, Canada) at a sampling rate of 2,343.75Hz. Participants sat upright in a dark, magnetically shielded chamber. Stimuli were projected onto a screen using either an LCD projector (Sanyo PLC-XP41, Moriguchi, Japan) or a DLP LED PROPixx projector (VPixx, Saint-Bruno, Canada) at 60 Hz refresh rate. Eye movements were tracked with an eyetracking system (Eyelink 1000, SR Research) at a sampling rate of 1000 Hz.

### Preprocessing

MEG Data was low-pass filtered at 10 Hz (two-pass forward-reverse Butterworth filter, order 4) and down-sampled to 20 Hz. We used robust detrending [25] to remove polynomial trends from the MEG data in a piece-wise fashion (600 s pieces, removal of linear trend followed by 10^th^ order polynomial). Individual noisy channels and trials were defined as those exceeding 10 times the standard deviation of the variability across channels or trials, respectively, and excluded from the analysis. For all temporally resolved analyses, results were smoothed using a 100 ms Hanning window (full width at half maximum).

### Data exclusion

Trials with blinks or eye movements outside of the fixation window were aborted and thus automatically excluded from analysis. Eye movements could not be measured for three participants, but their exclusion did not significantly change the results.

For some participants we were unable to analyse all 12 blocks due to technical issues. In one case this led to the exclusion of an entire coherence condition (high/full) due to a lack of counterbalancing across choice-response mappings.

Two participants were excluded due to insufficient behavioral performance caused by a misunderstanding of the task. Two participants were excluded due to technical problems with stimulus presentation or data recording. For reaction-time aligned choice analysis (Fig 3B), two participants were excluded due to technical issues.

### Source reconstruction

As MRIs for individual participants were not available, we source reconstructed the data using a standard MNI template brain. We generated a single-shell head model [26] and for each participant estimated three-dimensional (x, y, and z-direction) MEG source activity at 457 equally spaced locations 7 mm beneath the skull, using linear spatial filtering (beamforming) [27]. We retained, for each source, activity in all 3 directions. For the searchlight analysis, we used each of the 457 sources’ immediate neighbours, including all 3 dipole directions. We averaged data within 5 intervals with respect to stimulus onset (−0.85-0 s; 0-1 s; 1-2 s; 2-3 s; 3-4 s).

### Cross-validated MANOVA

We applied a cross-validated MANOVA (cvMANOVA) on the MEG data from single participants to estimate neural information about each of the variables of interest [19,28]. cvMANOVA constitutes an extension of the commonly used cross-validated Mahalanobis distance and allows for the simultaneous estimation of neural data variability due to several variables of interest. This estimation is performed in relation to unexplained noise variability. We therefore first estimate a baseline noise covariance matrix, using all trials from all possible unique combinations of variables or ‘conditions’. For each unique condition, beta weights are estimated and contrasted between conditions in cross-validation fold ‘training’ and ‘test’ sets separately. An estimate of true pattern distinctness is computed as the dot product of these contrasts, normalised by the noise covariance:

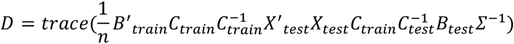

where X_test_ is the design matrix indicating the unique condition of each trial in the test set, C_train is_ the contrast vector the model is trained on, C_test_ the test contrast vector and ∑^-1^ the inverted noise covariance matrix. B_train_ and B_test_ contain the regression parameters of a multivariate general linear model:

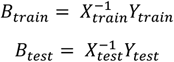

where Y_train_ and Y_test_ are the training and test datasets. The inverted noise covariance matrix was estimated from the mean activity during the time period starting from −0.85 seconds pre-stimulus to 2 seconds post-stimulus:

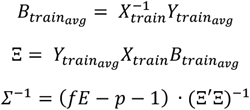

With fE being the degrees of freedom and p the number of sources used. Ξ was regularised towards the unity matrix using a regularisation parameter of 0.05.

Given that the design matrix and contrast vector include all unique conditions i.e. all possible combinations of variable levels, cvMANOVA independently quantifies information about each variable of interest, while not being confounded by information about the other, potentially correlated variables. In other words, cvMANOVA quantifies the pattern distinctness explained by each variable after discounting the patterns explained by all other variables included in the model. Importantly, cvMANOVA effectively controls imbalances in the distribution of trials over conditions without explicit stratification and the resulting loss of data.

Prior to cvMANOVA we reduced the dimensionality of the data using PCA. We computed a de-mixing matrix on the condition means of the training data only, and subsequently applied it to the test data. We selected the first 100 components for further analysis.

For all analyses we performed two-fold cross validation and 10 repetitions of cvMANOVA with different random seeds. We averaged results across repetitions and folds.

### Task variables and stimulus correction

To accurately estimate neural information about variables independently of each other, we had to ensure that all combinations of the variables of interest were present, including those under experimental control (coherence, choice-response mapping) and those dependent on the participants behaviour (choice, response). In all cvMANOVAs we included choice, response and coherence as variables.

As each stimulus pulse was randomly selected from 10 possible movies for each direction, there were many unique 2 s stimuli, such that the full stimulus could not be included as a variable. Given that stimulus and choice are correlated for non-chance performance, we used a stimulus correction procedure for those analyses assessing neural choice information independent of the stimulus (Figs 3, 4 and 5). For this, we computed the average neural activity for each pulse position, motion direction and coherence, and subtracted the 24 pulse-related averages from the neural data of individual trials based on the pulses presented on these trials. We then added back the pulse-related activity averaged across motion directions, independently for pulse position and coherence. This conservative correction likely removes choice-related neural activity that is strongly correlated with the stimulus. As this results in different over-corrections depending on performance, we did not interpret differences in choice information between coherence conditions.

To estimate the neural information associated with each individual pulse, we computed cvMANOVAs with the additional variable pulse direction, for each pulse position independently.

### Cross-variable information

To assess whether choice and response shared a common representational space, we measured cross-variable information. We implemented this by using a training contrast C_train_ differentiating between choice levels, and a test contrast C_test_ differentiating between response levels, and vice versa.

Notably a possible relationship between choice and response representations could vary in directionality across participants, such as for example an arbitrary choice-response associations that is unrelated to the actual choice-response mapping (e.g. an association between the left button and upwards motion). Thus, we did not average the resulting cross-variable information metric but compared absolute cross-variable information across participants (Fig 5).

Given that the maximal amount of shared information between two variables depends on the information available for each variable independently, it was important to take the strength of the individual representations into account. We therefore compared the measured cross-variable information to an estimate of the cross-variable information that could be expected for identical representations of variable strength [28]

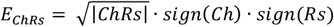

where Ch and Rs denote the pattern distinctness for choice and response, respectively, and E_ChRs_ is the expected cross-variable information. If representations were identical, the measured cross-variable information is expected to approach E_ChRs_, whereas cross-variable information small than E_ChRs_ indicates non-overlapping representations [28].

### Statistical analysis

We assessed the statistical significance of information using cluster-based permutation tests. After determining temporally contiguous clusters during which pattern distinctness was higher than 0 (one-tailed t-test over participants, p < 0.05), we randomly multiplied the information time-course of each participant 1,000 times with either 1 or −1. In each random permutation, we recomputed information clusters and determined the cluster mass of the strongest cluster. Each original cluster was assigned a p-value by comparing its mass to the distribution of masses of the random permutation’s strongest clusters.

To assess the cortical distribution of neural information we performed analogous cluster permutation tests, but across space and time.

To determine whether the multivariate patterns underlying choice and response were significantly different, we used one-tailed paired t-tests. We tested whether the absolute cross-variable information was smaller than the expected cross-variable information, and whether the absolute cross-variable information was higher than would be expected by chance. For the latter we ran cvMANOVA on data with shuffled choice labels and tested the absolute cross-variable information against this value. Tests were performed on the time period with robust choice and response information across participants (1.5-3 s).

### Software

Experimental code was written in MATLAB (Mathworks) using custom code and Psychophysics Toolbox extensions [29]. All analyses were performed in MATLAB using custom code as well as the Fieldtrip toolboxes [30].

## Acknowledgements

We thank Gabi Walker-Dietrich and Jürgen Dax for assistance with MEG recordings. This research was supported by the European Research Council (ERC; https://erc.europa.eu/) StG 335880 and CoG 864491 (M.S) and Deutsche Forschungsgemeinschaft (DFG; German Research Foundation; https://www.dfg.de/) project 276693517 (SFB 1233) (M.S.). The authors acknowledge support by the state of Baden-Württemberg through bwHPC, by the German Research Foundation (DFG) through grant no INST 39/963-1 FUGG (bwForCluster NEMO), and by the Open Access Publishing Fund of the University of Tübingen. The funders had no role in study design, data collection and analysis, decision to publish, or preparation of the manuscript.

## Author Contributions

Conceptualization: M.S., K.R.Q., N.N.; investigation: N.N., E.Z.; formal analysis: K.R.Q.; writing – original draft preparation: K.R.Q.; writing – review and editing: M.S., K.R.Q., F.S., N.N., E.Z.; supervision: M.S.; resources: M.S., F.S.; funding acquisition: M.S

## Competing Interests

All authors declare no competing interests.

## Notes

### Competing Interest Statement

The authors have declared no competing interest.

